# Microbiome remodeling through bacterial competition and host behavior enables rapid adaptation to environmental toxins

**DOI:** 10.1101/2023.06.21.545768

**Authors:** Dan Kim, Olga Maria Pérez-Carrascal, Catherin DeSousa, Da Kyung Jung, Seneca Bohley, Lila Wijaya, Kenneth Trang, Sarah Khoury, Michael Shapira

## Abstract

Human activity is altering the environment in a rapid pace, challenging the adaptive capacities of genetic variation within animal populations. Animals also harbor extensive gut microbiomes, which play diverse roles in host health and fitness and may help expanding host capabilities. The unprecedented scale of human usage of xenobiotics and contamination with environmental toxins describes one challenge against which bacteria with their immense biochemical diversity would be useful, by increasing detoxification capacities. To explore the potential of bacteria-assisted rapid adaptation, we used *Caenorhabditis elegans* worms harboring a defined microbiome, and neomycin as a model toxin, harmful for the worm host and neutralized to different extents by some microbiome members. Worms raised in the presence of neomycin showed delayed development and decreased survival but were protected when colonized by neomycin-resistant members of the microbiome. Two distinct mechanisms facilitated this protection: gut enrichment driven by altered bacterial competition for the strain best capable of modifying neomycin; and host avoidance behavior, which depended on the conserved JNK homolog KGB-1, enabling preference and acquisition of neomycin-protective bacteria. We further tested the consequences of adaptation, considering that enrichment for protective strains may represent dysbiosis. We found that neomycin-adapted gut microbiomes caused increased susceptibility to infection as well as an increase in gut lipid storage, suggesting metabolic remodeling. Our proof-of-concept experiments support the feasibility of bacteria-assisted host adaptation and suggest that it may be prevalent. The results also highlight trade-offs between toxin adaptation and other traits of fitness.

## Background

Animals harbor large gut microbial communities, microbiomes, that are important for their health and fitness. In vertebrates, the gut microbiome has been shown to contribute to processes as diverse as development, immunity, metabolism and even mood regulation ^1–4^. In herbivores (both vertebrate and invertebrates), gut bacteria are essential for mere survival, as energy harvest from plants depends on bacterial enzymes ^5–7^. While current understanding highlights how interwoven are the fates of animals and their gut bacteria, the full scale of this dependence is yet unknown. To consider some numbers, the human gut microbiome is estimated to consist of as many bacteria as there are cells in the body and to encode 100-fold more genes than there are in the human genome ^8^. Thus, it is likely that our current understanding of the contributions of the gut microbiome represents only the tip of an iceberg.

Host-adapted symbionts, such as those of sap-eating aphids, open new niches by providing their hosts with essential nutrients otherwise missing in available diet, or provide protection from local threats ^9^. Niche adaptation can be facilitated also by looser interactions with gut bacteria. Adaptation to toxins is a case in point. To deal with chemically-diverse toxins, animals possess several families of detoxifying enzymes with relatively broad substrate specificity, for example the cytochrome P450 superfamily ^10^. However, the repertoire of such enzymes is constrained by genome size. Bacteria, on the other hand, as a group offer a far greater biochemical diversity, which throughout animal evolution has been instrumental in enabling host adaptation to toxins, particularly to plant toxins. Examples include the coffee berry borer, a pest of the coffee industry, which is among the few insects that can withstand caffeine, thanks to the metabolizing activity of *Pseudomonas fulva* gut commensals; or the beetle *Psylliodes chrysocephala,* a major pest of oilseed rape, which is protected from plant isothiocyanates thanks to *Pantoea* commensals; or the Mojave desert woodrat, which can subsist on the otherwise toxic creosote bush, thanks to yet uncharacterized gut commensals ^11–13^.

The last century has exacerbated the challenge of environmental toxins, with an expansion in human-made chemicals, such as pesticides, flame retardants, or even antibiotics (of which many are synthetic or semi-synthetic). Some of the released chemicals have intended toxic effects on pathogens or pests but may also have off-target toxicity, others are thought to be toxic but data is ambiguous, and many others have not yet been characterized for possible toxic effects. Such environmental toxins represent novel paradigms of toxicity and stress for animals (including humans) as they contaminate large areas within a short time and spread through water ^14^. The time scale of such exposures further reduces the potential of genetic variation within animal populations to provide an answer. Bacteria are better suited to offer that. An example of bacteria-assisted rapid adaptation is presented by the bean bug *Riptortus pedestris,* a soybean pest, which was shown to develop resistance to sprayed organophosphate pesticides through rapid acquisition of pesticide-metabolizing *Burkholderia*, thanks to their increased environmental abundance in sprayed fields ^15, 16^. While this example supports the feasibility and flexibility of bacteria-assisted adaptation, how prevalent it is, and what are the underlying mechanisms is yet unknown.

Here, we describe proof-of-concept experiments, using a model host and a model toxin, to explore the likelihood and mechanisms of bacteria-assisted adaptation. As a host model we used the nematode *Caenorhabditis elegans*, a recently established model for microbiome research, shown to harbor a diverse yet characteristic gut microbiome shaped by environmental availability, host genetics and interbacterial interactions ^17–23^. As a convenient model toxin we chose neomycin, an aminoglycoside antibiotic that is also toxic to worms ^24^. As an antibiotic, neomycin is bound to affect the worm gut microbiome, while resistant community members could modify and neutralize it and help the host adapt. The experiments described below demonstrate that bacteria-assisted rapid adaptation could occur through distinct mechanisms in the bacterial community and in the host, leading to reproducible microbiome remodeling and enrichment for protective bacteria. The mechanistic redundancy and the reproducibility of this process increases the likelihood that bacteria-assisted adaptation is indeed prevalent, in alleviating environmental toxicity and possibly also in enabling adaptation to other types of environmental changes. At the same time, we considered that changes in microbiome composition also represented dysbiosis, which is often associated with pathology ^25, 26^. Exploring the long-term consequences of adaptive microbiome remodeling identified trade-offs between the short-term toxin adaptation and longer-term effects on infection resistance and on metabolic remodeling. Such trade-offs should be taken into account when considering the final outcomes of this adaptation.

## Results

### Gut bacteria protect *C. elegans* from neomycin toxicity

Worms were raised either on non-colonizing *E. coli* or on CeMbio, a community of twelve characterized worm gut commensals selected to represent the *C. elegans* core gut microbiome ^27, 28^. Analysis of neomycin sensitivity among CeMbio members revealed a wide range, including four strains, boxed in Fig. 1A, which maintained growth following neomycin exposure, both on plates and at least to some extent in liquid cultures. Worms raised on CeMbio were protected from neomycin toxicity. They completed normal development, unlike those raised on *E. coli* (Fig. 1B), and when shifted to neomycin as adults, survived significantly longer, demonstrating protection attributed to their gut bacteria (Fig. 1C). Together, these results demonstrate that the gut microbiome assembled from CeMbio can help its host adapt to neomycin.

**Figure 1.**
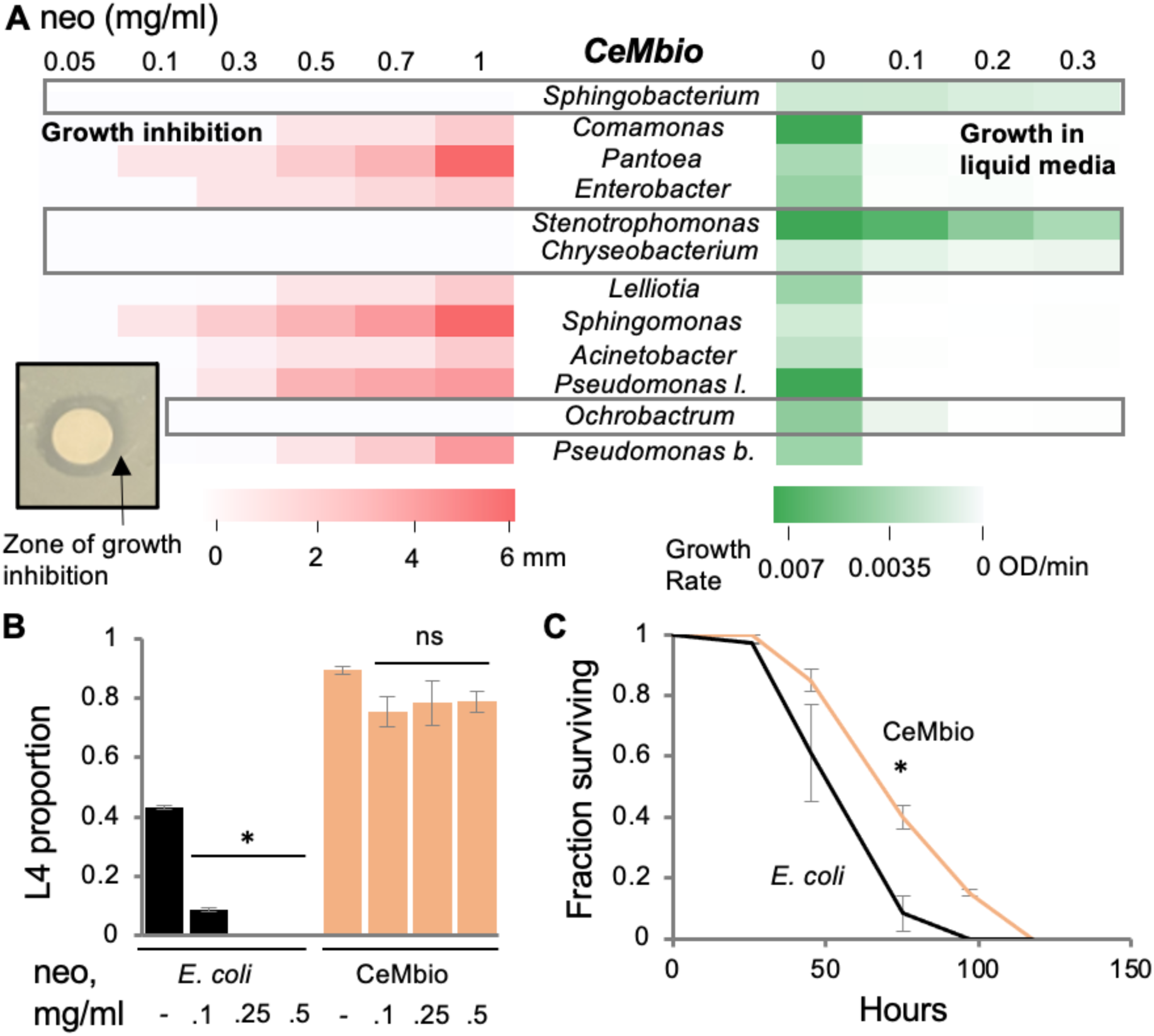
The gut microbiome protects host from neomycin toxicity. **A.** Sensitivity to neomycin, quantified in CeMbio community members by disk diffusion growth inhibition (red, see inset), or by liquid media growth (green). Resistant strains are boxed. **B.** Effects of neomycin on development of worms raised on *E. coli* or on CeMbio. Proportion of worms reaching L4 following 30 hours of growth from L1 at 20°C. Shown are averages ± SD for two plates, with 35-70 worms each, from a representative experiment out of two with similar results; *, p < 4E-4. **C.** Survival of wild type worms raised on designated bacterial cultures and shifted as L4 (t=0) to plates with 0.5 mg/ml neomycin. Shown are averages ± SD for two plates in a representative experiment out of two with similar results; N=54-77/group; *, p=2E-8, logrank test.

### Microbiome adaptation involves gut enrichment with protective *Stenotrophomonas* independent of the extent of its environmental availability

Worms raised without neomycin on monocultures of the four neomycin-resistant strains and shifted in early adulthood to plates with neomycin demonstrated that each of the resistant bacterial strains on their own could protect worms from subsequent exposure to neomycin, as opposed, for example, to *Lelliotia*, a moderately-sensitive strain, which could not, although it normally was an effective colonizer of worms (Fig. 2A) ^28^. This linked bacterial neomycin resistance with the ability to protect the host. However, next-generation sequencing of gut microbiomes showed that only one of the four – *Stenotrophomonas indicatrix* JUb19, took over the gut microbiome in worms exposed to neomycin (Fig. 2B and Supplementary Table 1). Whereas microbiome composition of worms raised without antibiotics showed enrichment for *Ochrobacterum* and *Enterobacteriacae* (*Lelliotia* and *Enterobacter* could not be distinguished based on their 16S V4 region) compared to the lawn environment, worms raised in the presence of neomycin showed enrichment for *Stenotrophomonas* (Fig. 2B). The pattern emerging in several similar experiments was that gut microbiomes in worms raised on CeMbio without antibiotics were dominated by *Ochrobactrum*, while gut microbiomes in worms exposed to neomycin were dominated by *Stenotrophomonas* (Fig. 2C left and Supplementary Table 1). The driving force for the *Stenotrophomonas* enrichment seemed to be distinct from its environmental availability, which changed relatively little in CeMbio lawns with or without neomycin (the latter probably because bacteria in lawns are not growing)(Fig. 2B). Additional experiments, in which worms were raised on CeMbio and were colonized without neomycin, then shifted to neomycin with *E. coli* as food, still showed enrichment of *Stenotrophomonas* indicating that the change in microbiome composition was due to the effect of neomycin on, or in, the worm gut (Fig. 2C right and Supplementary Table 1).

**Figure 2.**
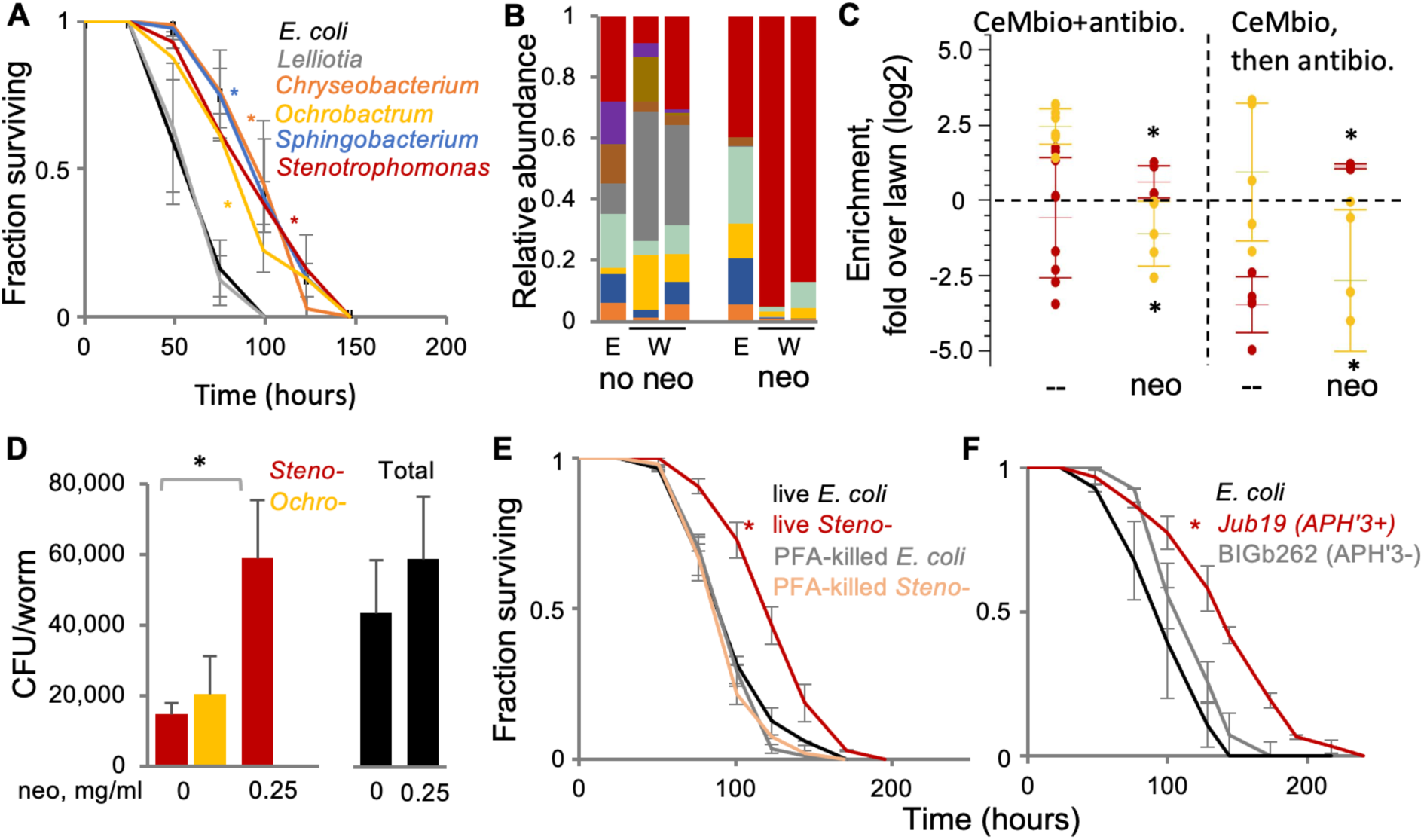
Neomycin exposure enrich for neo-resistant gut *Stenotrophomonas*. **A.** Survival assays of wild type worms raised on monocultures of designated CeMbio members until gravid and shifted to plates with 0.5 mg/ml neomycin (and *E. coli* as food). Shown are averages ± SD for one experiment performed in duplicates of two with similar results; N=62-82/group; *, p < 2E-08, logrank test. **B.** Microbiome composition of wild type worms raised on CeMbio +/-0.1 mg/ml neomycin (W, N=50/group) or in their lawn environment (E). Each bar graph represents an independent worm population, with relative abundance of strains colored as in A. **C.** Gut enrichment of *Ochrobactrum* and *Stenotrophomonas* in worms raised with +/-0.1 mg/m neomycin or kanamycin (left, n=3 independent experiments, with 2-3 separate populations each), or worms raised on CeMbio and shifted to antibiotics (with *E. coli* as food) as early gravid (n=2 independent experiments, 2-3 separate populations each). Shown are averages ± SD, *, p<0.05, t-test. **D.** Bacterial load in gravid worms (N=15/group) raised on CeMbio +/-neomycin. Shown are CFU counts on LB with 0.1 mg/ml neomycin (left) or without (right); averages ± SD for one experiment of four plates of two experiments with similar results. *, p=0.0018, t-test. **E, F.** Survival assays of wild type worms raised on monocultures of the designated *bacteria (*or dead, when designated) and transferred as gravid to plates with 0.25 mg/ml neomycin (and *E. coli* as food). Shown are averages ± SD for two plates, N=57-83/group *, p<7E-7, logrank test.

Experiments similar to those described above of worms raised on CeMbio with or without neomycin, but using colony forming units (CFU) counts to estimate the actual number of bacterial cells in the worm gut demonstrated that gut *Stenotrophomonas* enrichment was not only an increase in its relative abundance, but also in the absolute number of *Stenotrophomonas* cells per worm (Fig. 2D). This result suggested that gut abundance of *Stenotrophomonas* increased thanks to its release from competition, probably with *Ochrobactrum*, the otherwise dominant strain. *In vitro* competition experiments in liquid culture, showing a similar increase in *Stenotrophomonas* growth and displacement of *Ochrobactrum* following addition of neomycin (Supplementary Fig. 1), support the hypothesis that the *Stenotrophomonas* increase is due to release from competition with *Ochrobactrum*.

Scanning the genome of the JUb19 *Stenotrophomonas* strain for potential aminoglycoside-protective mechanisms identified an aminoglycoside 3’-phosphotransferase homolog, APH(3’), which in other bacteria was shown to phosphorylate and neutralize aminoglycosides such as neomycin and kanamycin ^29^. The encoded enzyme may be the factor responsible for Jub19’s neomycin resistance and for its ability to protect the host. Worms raised on *Stenotrophomonas* killed with paraformaldehyde were as sensitive to neomycin as those raised on *E. coli*, indicating that active bacterial protein expression is required for protection (Fig. 2E). Scanning the genomes of several other *Stenotrophomonas* species (gratefully received from Michael Herman, University of Nebraska), we identified BIGb262, a *Stenotrophomonas rhizophila* strain, also isolated form *C. elegans*, which shared 80.8% of its nucleotide sequence with Jub19 (H. Schulenburg, personal communication), but lacked the APH(3’) gene. Unlike Jub19, BIGb262 was unable to protect worms from neomycin, providing support (albeit correlational) for the putative role of Jub19’s APH(3’) in neomycin protection (Fig. 2F).

### *kgb-1*-dependent toxin avoidance enables preference and acquisition of neomycin-protective strains

The results described above demonstrate that exposure to neomycin reshaped the interactions between gut bacteria, resulting in enrichment of APH(3’)-positive *Stenotrophomonas*. In these experiments, *Stenotrophomonas* was made available to worms mixed with all other members of CeMbio. However, in nature, it is plausible that worms encounter bacteria in spatially segregated compartments where they proliferate locally and form colonies. Furthermore, previous work from our lab has demonstrated that worms can prefer beneficial bacteria over non-beneficial congenerics ^30^. Therefore, we examined whether worms exposed to neomycin would gravitate toward neomycin-protective bacteria. Indeed, when worms exposed to neomycin were allowed to choose between CeMbio and *E. coli*, they significantly preferred CeMbio (Fig. 3A).

**Figure 3.**
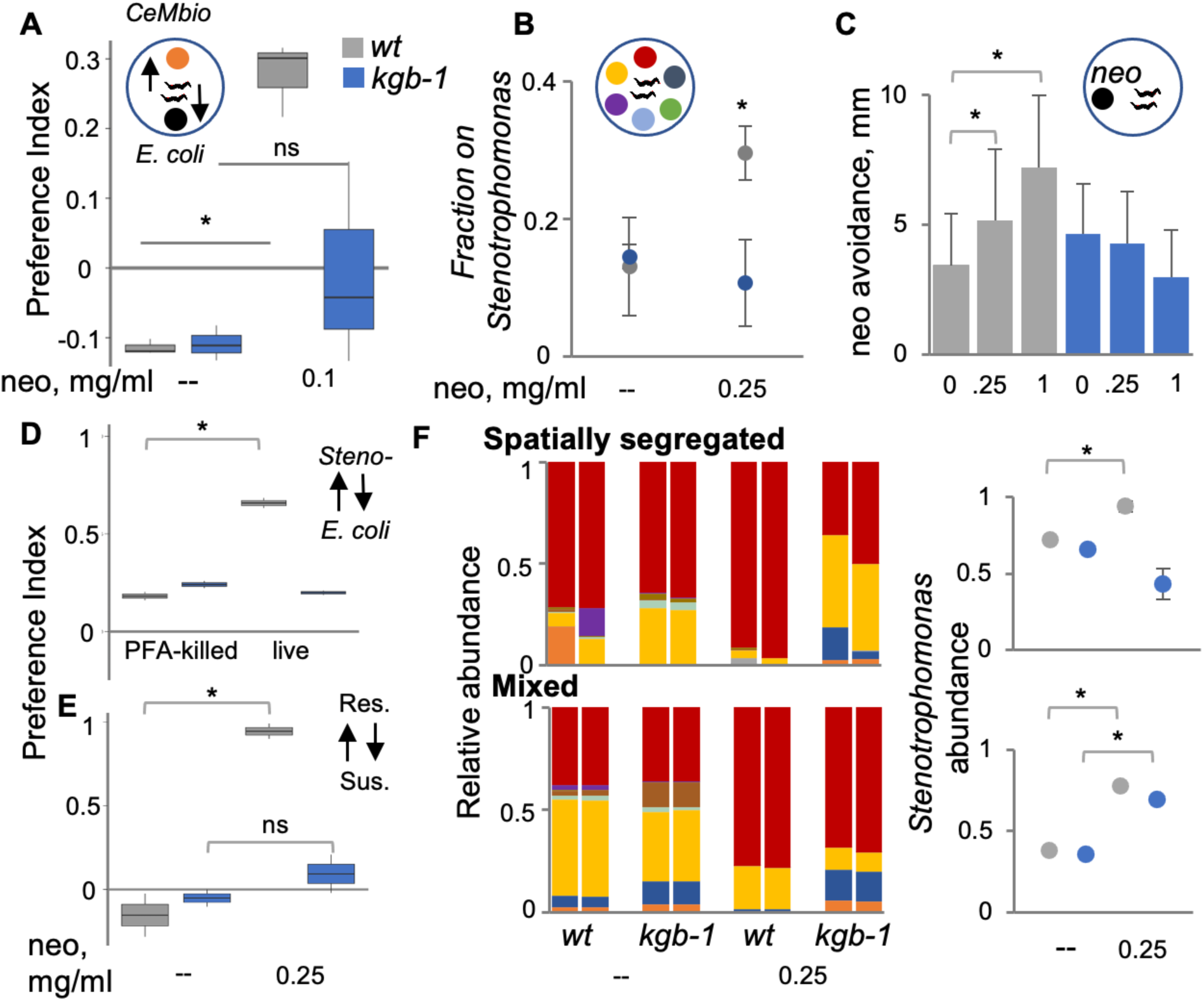
*kgb-1*-dependent toxin avoidance enables preference and acquisition of neo-protective strains. **A.** Preference assay of wild type or *kgb-1* worms (N=100/group) between CeMbio and *E. coli* with or without neomycin. Shown are median (lines) and interquartile (boxes) performed in triplicates; *, p=0.0002. A representative experiment of two with similar results. **B.** Preference assay of wild type worms in a multi-choice assay on spatially segregated monocultures of CeMbio members, with or without neomycin. Shown are averages ± SD for two plates (n=46-185/plate), *, p=0.0046. **C.** Avoidance of neomycin in wild type or *kgb-1* gravid worms evaluated based on worm distance made in one hour from the point of neomycin application. averages ± SD for two plates; N=75/group, *, p<2 E-7. **D,E.** Pairwise preference assays of wild type or *kgb-1* worms between *E. coli* or *Stenotrophomonas* (Jub19), live or dead (E), or between resistant *S. indicatrix* (Jub19) and susceptible *S. rhizophila (*BIGb262). Shown are median and interquartile of one experiment performed in duplicates of two with similar results, N=54-157/group, *, p<0.02. **F.** Left, gut microbiome composition in worms raised on spatially segregated cultures of CeMBio members as in B, or on a mix of all CeMbio members, with or without neomycin. Colors, as in Fig. 1 (red, *Stenotrophomonas*, yellow, *Ochrobactrum*). Right, *Stenotrophomonas* abundance in experiments presented on the left panel. averages ± SD for two populations, N=80/group; *, p<0.05, t-test.

Previous studies have demonstrated a role for KGB-1/JNK, a stress-activated MAP kinase, in initiating an avoidance behavior leading worms away from conditions where essential cellular processes were inhibited ^31^. The effects of neomycin on worms – developmental arrest or death, indicated inhibition of similarly essential (though yet unknown) processes. This suggested that KGB-1 might be involved also in neomycin-induced preferences. Examination of *kgb-1* mutants provided support for this hypothesis, demonstrating that *kgb-1* mutants exposed to neomycin did not show preference for the beneficial CeMbio over *E. coli*. A similar trend was observed when worms were allowed to choose between the different members of CeMbio, in this case spatially segregated. Higher percentage of wildtype animals exposed to neomycin was found on the *Stenotrophomonas* colony, but not so *kgb-1* mutants (Fig. 3B). The involvement of KGB-1 further suggested that worms showing preference for neomycin-protective bacteria may be driven by avoidance of active toxin. To test whether neomycin induced avoidance, we applied neomycin in different concentrations to worm plates and measured the distance of individual worms from the point of application (Fig. 3C). This demonstrated concentration-dependent avoidance, which was *kgb-1*-dependent, supporting a role for KGB-1 in responding to the toxic effects of neomycin and activating a behavioral response that led worms away from the toxin, and as a consequence, toward toxin-neutralizing bacteria where concentrations were lower. In agreement with the proposed mechanism, *kgb-1*-dependent preference was only of live *Stenotrophomonas* (Fig. 3D) and was stronger for the APH(3’)-expressing Jub19 (Fig. 3E). Interestingly, the three non-enriched neomycin-protective CeMbio members (Myb71, BIGb0170 and JuB44) were also preferred by worms exposed to neomycin in pairwise assays against *E. coli* (Supplementary Fig 2). This suggested that in spite of not having a recognizable neomycin-modifying enzyme, these strains might still be able to modify and neutralize neomycin (which also explains their ability as monocultures to protect worms from neomycin (Fig. 2A)).

To test how *kgb-1*-dependent preferences could affect gut microbiome composition, we raised wildtype and *kgb-1* worms with or without neomycin, and on *E. coli* or CeMbio – the latter either with all strains mixed, as in previous experiments, or spatially segregated. In the configuration in which worms could choose between different CeMbio strains, wildtype worms exposed to neomycin showed gut enrichment for *Stenotrophomonas*, but *kgb-1* mutants did not (Fig. 3F and Supplementary Table 1), indicating that *kgb-1*-dependent preference of *Stenotrophomonas* contributed to its gut enrichment. When CeMbio strains were mixed, both wildtype and *kgb-1* animals showed neomycin-induced *Stenotrophomonas* enrichment, indicating that when preference cannot play a role, *Stenotrophomonas* still became enriched, but this was independent of *kgb-1*. KGB-1 was previously shown to regulate transcriptional stress responses, including in the gut ^32^, but the results presented here indicate that KGB-1’s contribution to neomycin-induced changes in microbiome composition were not associated with its regulatory function in the intestine, but with its regulation of worm behavior. In summary, these results describe a *kgb-1*-dependent mechanism that drives worms to seek protective bacteria, and a second, distinct, *kgb-1*-independent mechanism that makes the gut niche more favorable to *Stenotrophomonas*.

### Long-term consequences of neomycin-induced changes in gut microbiome composition

Enrichment of the neomycin-resistant *Stenotrophomonas* in the gut protects worms from toxicity. However, this deviates from baseline gut microbiome composition and might represent gut dysbiosis. Therefore, it was of interest to find how neomycin-adaptive changes affected other features of host function and fitness. Wildtype worms were raised either on CeMbio, CeMbio with neomycin - to recreate the dysbiotic gut microbiome (with enrichment of *Stenotrophomonas*), and on a CeMbio subset reconfigured to emulate the composition of the gut microbiome in worms exposed to neomycin, but without the toxin itself. CFU counts of gut microbiomes in worms raised on this reconfigured community demonstrated the desired enrichment for *Stenotrophomonas*, although not to the same extent as in neomycin-treated worms (Supplementary Fig. 3). Starting with major life history traits, we found no significant effect of the dysbiotic microbiome on lifespan, development rate and fecundity (not shown). However, a significant increase in gut lipid levels was observed in worms raised on *Stenotrophomonas*-enriched communities, representing an increase in energy storage (Fig. 4A). A small increase was observed in worms raised on CeMbio and exposed to neomycin, but an even larger increase was observed in worms raised on the reconfigured CeMbio without neomycin, and a similar increase was observed in worms raised on *Stenotrophomonas* alone, indicating that *Stenotrophomonas* enrichment was sufficient to alter lipid storage. Resistance to the pathogen *Pseudomonas aeruginosa* was also affected in *Stenotrophomonas*-enriched worms, showing a significantly lower resistance in worms raised on reconfigured CeMbio communities compared to that of worms raised on standard CeMbio (Fig. 4B). To test whether the decrease in infection resistance was due to a decrease in community diversity, or alternatively, the loss of specific protective strains, we generated three additional reconfigurations of the CeMbio community – one that included only strains that were previously shown to prevent pathogenic colonization of *P. aeruginosa* – *Sphingobacterium sp.* BIGb0170*, Chryseobacterium sp.* JUb44, and *Pantoea sp.* BIGb0393 (Supplementary Fig. 4) ^30^, a second, which lacked only these protective strains, and a third, in which three non-protective strains were excluded, reducing general diversity, but keeping the three protective strains. The results shown in Fig. 4C suggest that loss of infection-protective strains due to neomycin sensitivity is likely the reason for reduced infection resistance following neomycin-driven reconfiguration of the gut microbiome, not a general decrease in microbiome diversity.

**Figure 4.**
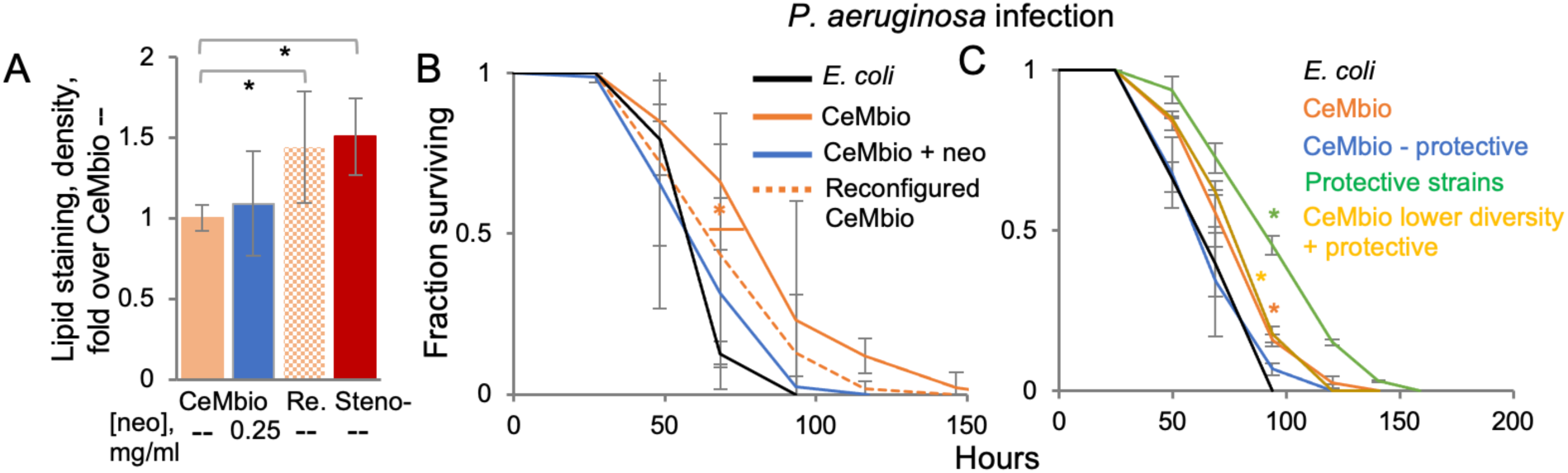
Consequences of neomycin adaptation. **A.** Lipid staining of wild type gravid worms (n=16-21/group) raised on bacterial communities as designated: CeMbio with or without neo; Re, reconfigured CeMbio; Steno-, *Stenotrophomonas* only. Shown are averages ± SD for one experiment of two with similar results, *, p<3E-5, t-test. **B, C.** Infection resistance of wild type worms raised on bacterial lawns as designated and shifted in adulthood to *Pseudomonas aeruginosa*. Protective strains include *Pantoea, Sphingobacterium,* and *Chryseobacterium.* Shown are averages ± SD for one experiment, performed in duplicates, of two experiments with similar results; N=42-74/group, *, p=0.004 and p<0.003 respectively, logrank test.

## Discussion

Using neomycin as a convenient model for environmental toxins - toxic to the *C. elegans* host and effectively altering the gut microbiome, our results demonstrate that the gut microbiome can provide protection from toxicity. In the context of a full community, protection was conferred by the most resistant member of the community, a *Stenotrophomonas indicatrix* strain, which persisted in the affected worm gut and took over the gut microbiome. In addition, exposed worms sought relief from the toxin, driven by avoidance behavior that depended on the stress-activated JNK homolog gene *kgb-1,* and that when given a choice resulted in preference for protective bacteria. Thus, two distinct mechanisms converge to facilitate colonization by protective bacteria. In this toxin-adapted microbiome, the enrichment for neo-resistant strains (primarily *Stenotrophomonas*) described a significant deviation from the baseline gut microbiome composition, representing dysbiosis. The effects of this dysbiosis on host fitness were limited, leaving development rate, fecundity and lifespan largely unaffected. However, it did increase the susceptibility of hosts to the pathogen *Pseudomonas aeruginosa*, driven by loss of pathogen-protective but neomycin-sensitive strains, and further altered metabolism leading to increased lipid storage. Our results demonstrate the feasibility of microbiome-dependent host adaptation to environmental toxins. They further demonstrate that this adaptation could be facilitated through more than one mechanism, either in bacteria or in the host, and that the mechanisms involved are of general purpose - interbacterial competition and avoidance of harmful environments, together increasing the likelihood that adaptation will occur. Our results focus on adaptation to an environmental toxin, but similar mechanisms could give rise to adaptation to other types of environmental stress. At the same time, our results highlight the trade-off that may take place between short-term adaptation to the toxin and other adaptive traits, some of which with potential long-term consequences.

While bacterial environmental availability can be affected by neomycin toxicity, environmental availability of the protective strain was not the main factor responsible for gut microbiome enrichment with *Stenotrophomonas*, as this occurred also in worms first colonized by CeMbio members and only subsequently exposed to the toxin. The increased bacterial load observed in the toxin-adapted worms is in line with a release from competition and suggests that the mechanism through which *Stenotrophomonas* becomes enriched is decreased competition or competitive exclusion of less-resistant gut bacteria. While previous studies of the bean bug adaptation to pesticide linked host resistance to acquisition of environmentally-enriched protective bacteria ^16^, our results demonstrate that when biochemical capabilities pre-exist in the gut microbiome, toxin-induced stress could be sufficient for microbiome remodeling and host adaptation, independent of environmental availability.

The choice of neomycin as the model toxin and concentrations selected for testing was to ensure microbiome changes in a community with somewhat limited diversity and to ensure observable effects on hosts. In the environment, antibiotic concentrations are usually far lower than those used here ^33, 34^. Nevertheless, there are examples of antibiotics, specifically aminoglycosides, used in the environment in concentrations as high as those we used ^35^. Thus, the mechanisms that we identified as taking part in worm adaptation to neomycin might play similar roles in natural contexts. These mechanisms are of general purpose. On the bacterial side, aminoglycoside modifying enzymes are common among bacteria, including in the human gut ^36–39^. On the host side, the *C. elegans* stress-activated MAP kinase KGB-1, a JNK homolog, is a conserved protein involved in diverse stress responses as well as in behavioral modulation ^40–42^. Together, such mechanisms could facilitate bacteria-assisted adaptation to toxic antibiotics in nematodes in the wild, as well as in other animals. Adaptation to other toxins may rely on different bacterial toxin-modifying enzymes but could be similarly feasible. And when appropriate enzymes are not as common as those involved in antibiotic resistance, environmental exposure to the toxin could significantly increase availability of bacteria expressing the appropriate enzyme, as found for the pesticide-resistance bean bug discussed earlier ^16^.

While remodeling of the worm gut microbiome provided relief from neomycin toxicity, some trade-offs were observed, i.e. increased lipid storage and susceptibility to infection. Maintaining gut microbiome composition within certain boundaries (still ill-defined) is essential for maintaining gut homeostasis ^43^. Deviations, often observed upon disruption of immune signaling, lead to dysbiosis and pathology ^22, 44, 45^. The *Stenotrophomonas* enrichment in neomycin-affected worms represents a significant deviation from the typical composition of the gut microbiome assembled from CeMbio and is sufficient to cause the observed increase in lipid storage, even without neomycin exposure. A previous study reported higher lipid storage upon worm infection with the opportunistic pathogen *Stenotrophomonas maltophilia* ^46^. However, our results with the non-pathogenic *P. indicatrix* suggest that metabolic remodeling might be distinct from the pathogenicity of *S. maltophilia*, and potentially associated with other interactions between *C. elegans* and members of the *Stenotrophomonas* genus. Host metabolism seems to be sensitive to changes in microbiome composition ^47^. The effects of *Stenotrophomonas* enrichment are but one example, but added to previous reports of gut-dysbiosis-induced metabolic remodeling, i.e. metabolic syndrome and obesity ^44, 48^, it may suggest that metabolic remodeling might be a common consequence of microbiome-assisted adaptation to toxins.

The results presented here support the notion that microbiome-assisted host adaptation to environmental toxins is straightforward, can be achieved through distinct routes and as a consequence, is probably more prevalent than currently appreciated. Increases in the spread of environmental toxins are only one facet of global change, perhaps representing a change in which bacteria are particularly likely to play a role. However, gut bacteria could contribute to any aspect of ecological adaptation. The microbiome was previously proposed as a potential source for adaptive novelty ^49^. It seems that helping hosts adapt to a changing environment may be one such contribution. It might be happening all around us, and may further be a yet unaccounted cause of health issues stemming from trade-offs with such adaptation.

## Methods

### Strains and reagents

*C. elegans* strains used in this study include the N2 wild-type strain and the mutant strain *kgb-1* (km21), both obtained from the *Caenorhabditis* Genetic Center. *E. coli* strain OP50, similarly obtained, was used as food and as control. BIGb262, a *Stenotrophomonas rhizophila* strain, was gratefully received from Michael Herman from the University of Nebraska (NCBI accession PRJNA986126). A GFP-expressing *Pseudomonas aeruginosa* derivative (PA14-GFP) was used in following bacterial accumulation as well as for infection resistance assays. Neomycin was purchased from Sigma (cat. No. N1876).

### The CeMbio community

CeMbio is a defined community of twelve characterized and genome-sequenced worm gut commensals ^28^, which include: CEent1 (*Enterobacter hormaechei*), JUb66 (*Lelliottia amnigena*), MYb10 (*Acinetobacter guillouiae*)*, JUb134* (*Sphingomonas molluscorum*), *JUb19 (Stenotrophomonas indicatrix*)*, MYb11* (*Pseudomonas lurida*)*, MSPm1 (Pseudomonas berkeleyensis), BIGb0172 (Comamonas piscis), BIGb0393 (Pantoea nemavictus), MYb71 (Ochrobactrum vermis), BIGb0170 (Sphingobacterium multivorum), and JUb44 (Chryseobacterium scophthalmum)*. In setting for an experiment, individual strains were grown at 28°C for two days, shaking, in 2 mL of LB. Saturated cultures of each monoculture were adjusted to OD_600nm_ of 4, mixed in desired combinations (or kept as individual strains) and spread on nematode growth media (NGM) plates with or without antibiotics.

A bacterial community configured to mimic the gut community as assembled from CeMbio in worms exposed to neomycin, included *Jub19, Myb71, and Bigb0172* in a ratio of 9: 0.27: 0.5, respectively.

### Worm populations

Germ-free and synchronized populations of arrested L1 larvae were obtained by bleaching gravid worms to release eggs, hatching eggs on NGM without food, and then raising worms to adulthood at 20°C on plates with bacterial monocultures or communities.

### Assessing bacterial susceptibility to antibiotics

#### Growth inhibition on plates

Sterile paper disks soaked in solutions with different antibiotic concentrations were placed on bacterial lawns seeded from 0.05 OD_600nm_ cultures on LB plates and incubated overnight at 28°C. Bacterial sensitivity to antibiotic was assessed by the width of the clear halo around the disk.

#### Growth in liquid

Individual strains, diluted from saturated cultures to 0.05 OD_600nm_, were raised in LB at 28°C in wells of a microtiter plate containing different antibiotic concentrations, and their growth monitored during 48 hours with OD_600nm_ readings every 5 minutes using a Molecular Devices Versamax plate reader. Maximal growth rate was calculated to assess antibiotic sensitivity.

#### *C. elegans* survival on neomycin

Worms were raised on OP50, CeMbio, or individual CeMbio strains until gravid, washed off plates and washed three times with 15ml filter-sterilized M9 before transferring to NGM plates with neomycin, and with *E. coli* as food, on which survival was scored. In experiments testing the ability of dead bacteria to protect worms from neomycin, killing was achieved by a 45-minute incubation at 37°C with 1% paraformaldehyde, followed by washing five times with 15 ml filter-sterilized M9.

### Genome sequence analyses

Scanning of bacterial genomes for potential antibiotic resistance genes was performed using the Comprehensive Antibiotic Resistance Database (CARD, https://card.mcmaster.ca). Genome sequences of CeMbio strains are available in the European Nucleotide Archive (accession number PRJEB37895). Genome sequence of BIGb262 was provided by Michael Herman, from the University of Nebraska.

### Harvesting and processing worms for colony forming units (CFUs) and next generation sequencing

Worms raised on CeMbio were washed off plates, and surface sterilized with bleach (following paralysis in 25 mM levamisole), as previously described ^28^, and the worms (15 worms in 250μl of M9) were ground using zirconia beads releasing live bacteria. After serial dilution, aliquots were plated onto LB plates +/-neomycin at 28°C for 2-3 days. For next generation sequencing, the washed worms were used for DNA extraction using the DNeasy PowerSoil Kit (#12888). DNA was similarly extracted from swabs taken of the bacterial lawns.

### Preparation of sequencing libraries

Extracted DNA was used as template (> 1 ng per reaction) for amplification of the 16S rDNA V4 region, using tailed 515f and 806r primers (see below; sequences in the bracket align to V4 region), compatible with the Nextera XT DNA library prep kit (Illumina, FC-131-2001), and KAPA HiFi HotStart polymerase (Roche, #07958935001). Amplification was carried out with 95°C for 3min, 25 cycles of 95°C for 30sec, 55°C for 30sec and 72°C for 30sec, and 72°C for 5min. Indices were added using the Nextera kit, according to the manufacturer instructions and products cleaned using AMpure XP reagent (Beckman Coulter, A63881). Libraries prepared from different samples were combined in equimolar ratios as suggested by the Illumina manual and used in paired-end next generation sequencing (NGS) using an Illumina MiniSeq.

515f: TCGTCGGCAGCGTCAGATGTGTATAAGAGACAG-[GTGCCAGCMGCCGCGGTAA]

806r: GTCTCGTGGGCTCGGAGATGTGTATAAGAGACAG-[GGACTACHVGGGTWTCTAAT]

### NGS data analysis

Analysis of 16S amplicon data was conducted using QIIME2 ^50^. In total, 96.9% of the total reads passed quality filtering with an average read of 81200 reads per sample. Sequences were aligned and clustered into operational taxonomic units (OTU) based on the closed reference OTU picking algorithm using the QIIME2 implementation of UCLUST ^51^ and the taxonomy of each OTU was assigned based on 99% similarity to reference sequences based on 16S sequences, available in the *Caenorhabditis* Genetics Center, of CeMbio strains.

### CFU counts

CFUs of *Ochrobactrum* and *Stenotrophomonas* were counted on LB plates with 0.1mg/ml Neomycin based on their distinct morphologies; small and dark colonies for *Ochrobactrum*, large and orangish for *Stenotrophomonas* (Supplementary Fig. S1). Estimates for total bacterial load relied on CFU counts on LB plates without antibiotics.

### Bacterial preference assays

Different bacterial strains/communities were seeded on the opposite sides of a 60mm NGM plate in pairwise choice assays, or around plates periphery in multiple choice assays. Synchronized L1 worms were transferred to the center of plates and the number of worms (#) in the vicinity of bacterial strains/communities was noted in each day until worms were gravid, and used for calculating a preference index: (# of worms on strain A – # on strain B) / total # of worms.

### Neomycin avoidance

30μl of neomycin solution in designated concentrations were spotted on a NGM plate with gravid worms and with OP50 as food. Images were captured with a Leica MZ16 F stereoscope equipped with a Qimaging Micropublisher 5.0 camera one hour later and the distance of worms from the point of antibiotic application was measured using ImageJ.

### Infection resistance assay

Worms raised on monocultures or bacterial communities until gravid were washed off and washed with 15ml filter-sterilized M9 solution three times, transferred to slow killing plates (SKP) with PA14, and survival was followed as previously described ^52^. Differences were assessed using Kaplan-Meier estimator and post-hoc logrank tests.

### Lipid quantification

Gravid worms raised on different bacterial strains/communities were washed off plates, washed three times in PBST (PBS + 0.01% Triton X-100 (Fisher Scientific, 9002-93-1), fixed in 40% isopropanol by three-minute incubation at room temperature and incubated with 3 mg/ml Oil Red O (Sigma, 09755) for 2 hours at room temperature, followed by two washes, 30 minutes each, with PBST ^53^. Images of stained worms were captured and signal levels quantified using ImageJ, normalizing to the unstained head region of worms.

## Acknowledgements

We thank Dr. Michael Herman and members of his lab in University of Nebraska for providing *Stenotrophomonas* strains, Dr. Britt Koskella - for the use of her lab’s microplate reader, Dr. Denis Titov - for useful comments and ideas, and Dr. Molly Matty - for help with lipid quantification protocol. This work was supported by NIH grants R01OD024780 and R01ES034012. DK was supported by National Science Foundation Graduate Research Fellowship Program (DGE 2146752).

## Author contributions

D.K. and M.S. conceived the project; D.K., assisted by S.B., L.W., D.J., K.T. and S.T. performed all experiments and analyzed their results, expect for experiments characterizing infection resistance to *Pseudomonas Aeruginosa*, which were carried out by O.P-C. and lipid staining experiments, which were performed and analyzed by C.D., D.K. and M.S. analyzed all experiments and together wrote the manuscript.

---

**Supplementary Figure 1.**
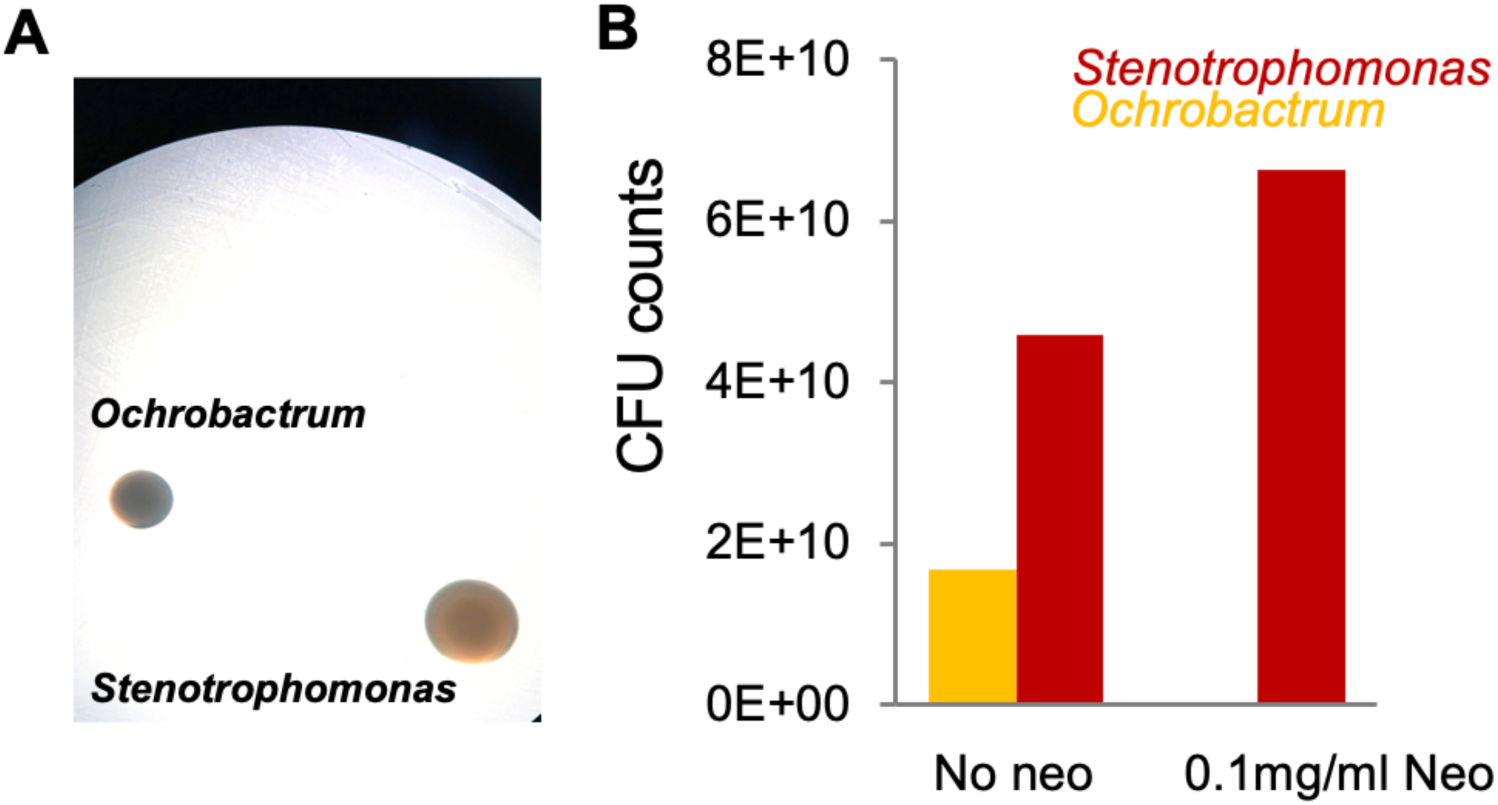
Release from competition with *Ochrobactrum* allows increased growth of *Stenotrophomonas.* **A.** Distinguishing between colonies of *Ochrobactrum* and *Stenotrophomonas* on LB with 0.1 mg/ml neomycin (48hours at 28°C). **B.** CFUs of designated bacteria, counted following co-culturing at 28°C for 48 hours in liquid LB +/-neomycin.

**Supplementary Figure 2.**
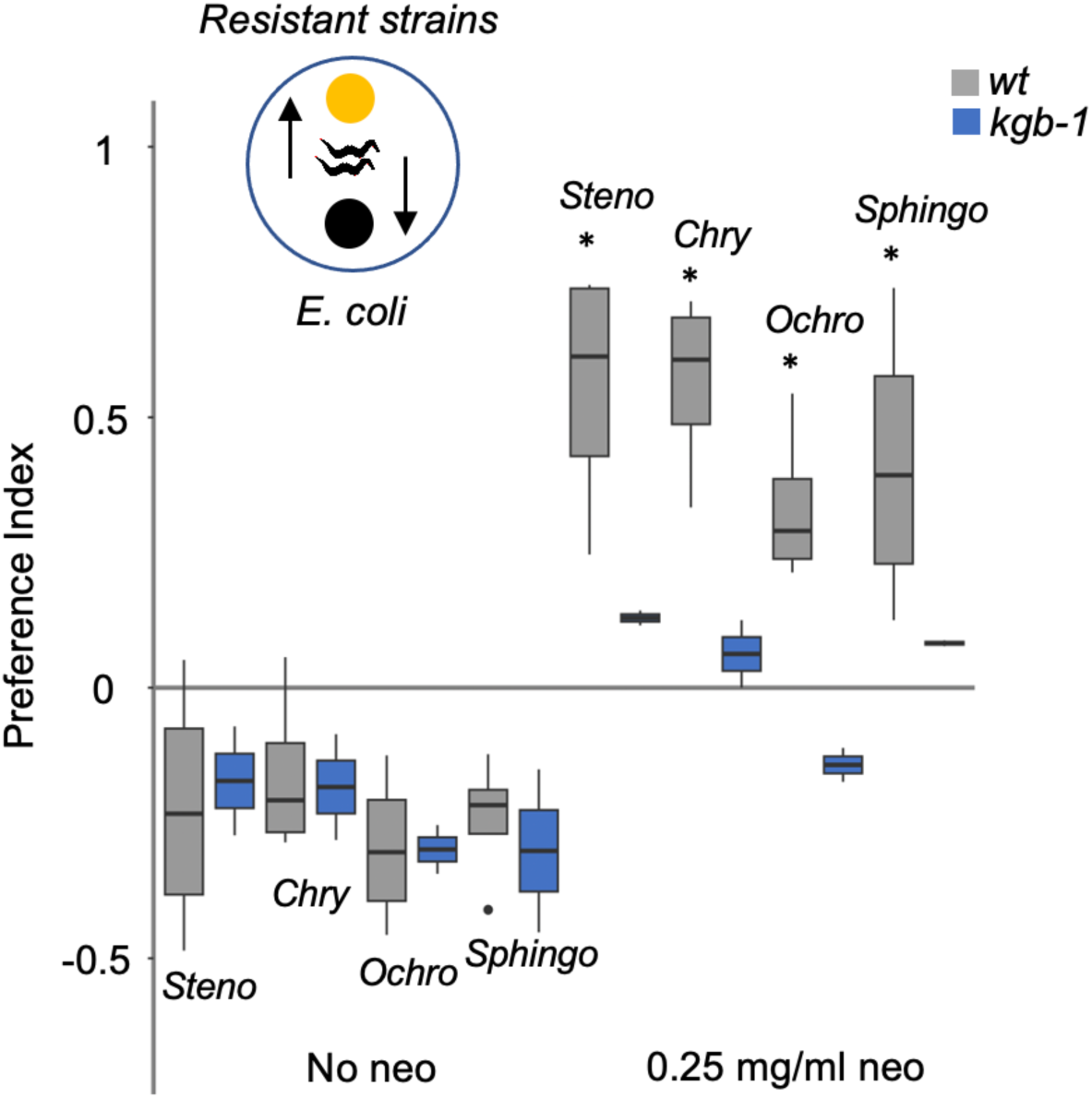
Neomycin and kgb-1-dependent preference for resistant CeMbio members. Preference assays of wild type or *kgb-1* worms between resistant strains, *Stenotrophomonas (Steno), Chryseobacterium (Chry), Ochrobactrum (Ochro),* and *Sphingobacterium (Sphingo)* (represented by yellow dot in scheme) and *E. coli*. Shown are median values (line) and interquartile (boxes) of one experiment performed in duplicates of two with similar results, n= 42-121 worms/group. *, p< 0.005, in comparison to respective no antibiotic exposure, t-test.

**Supplementary Figure 3.**
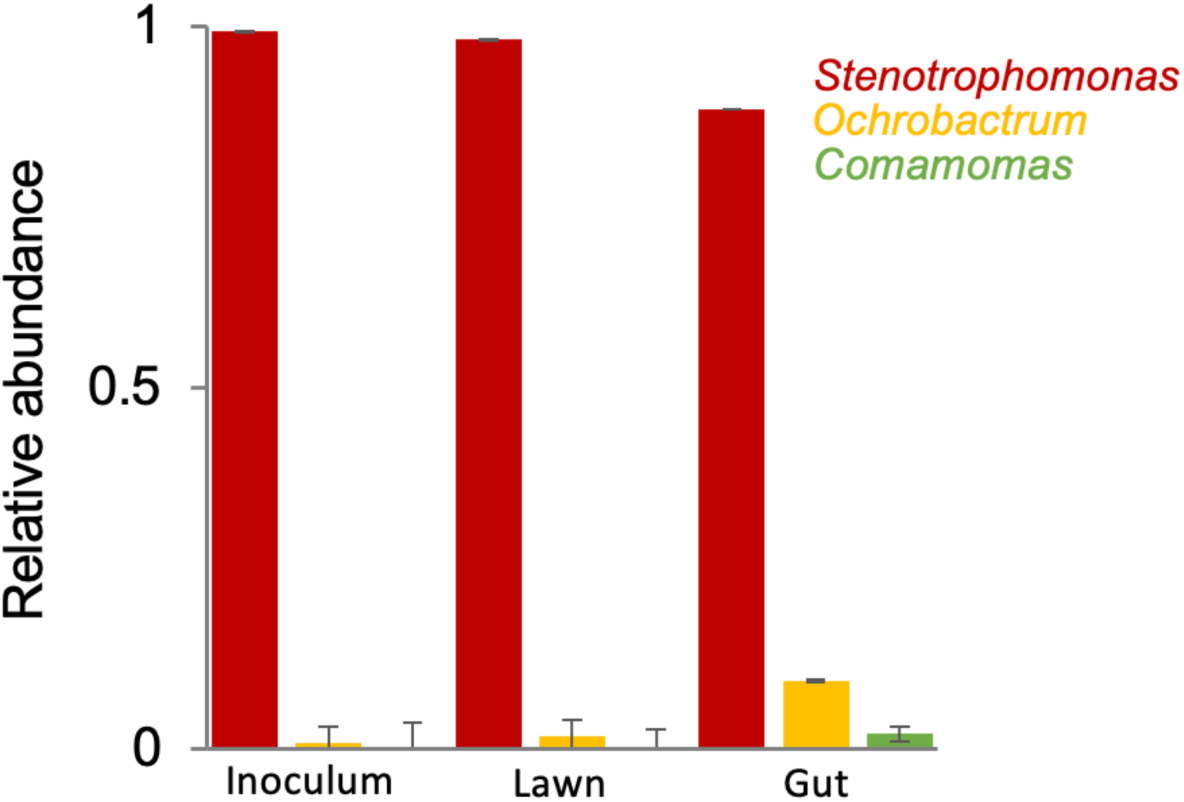
Evaluation of strain abundance in lawn and in worms following growth on the reconfigured CeMbio community. Relative abundance of the colored CeMbio strains in a lawn community emulating microbiome composition as in worms exposed to neomycin, and in worms raised on that lawn. Relative abundance was assessed based on CFU counts of respective strains on LB plates, distinguished based on colony morphology. Shown are averages ± SD for one experiment, performed in two plates, of two experiments with similar results for N=10 worms/group (Gut).

**Supplementary Figure 4.**
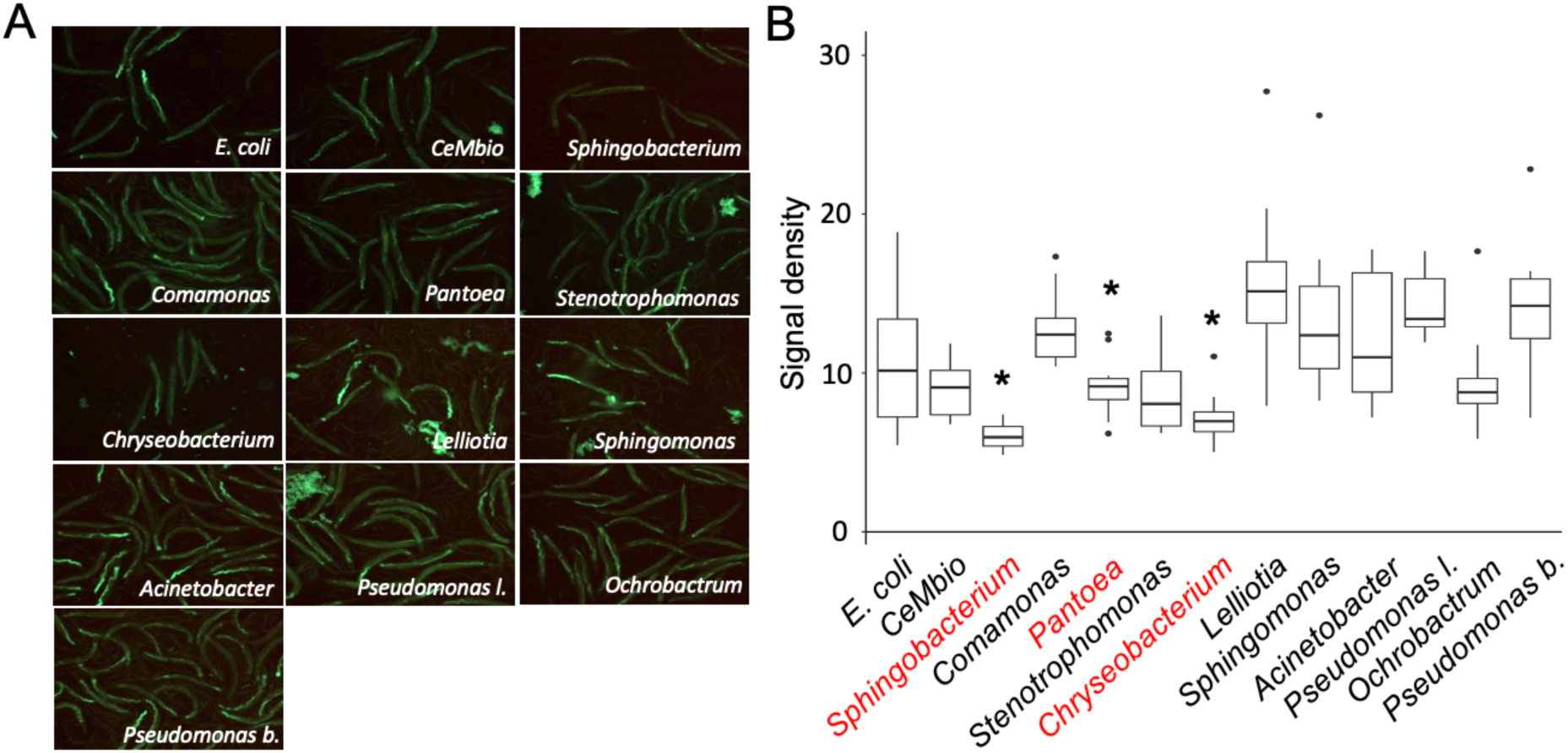
Differential ability of CeMbio strains to attenuate pathogenic colonization by *Pseudomonas Aeruginosa*. **A.** GFP fluorescence images of wild type *w*orms raised on monocultures of CeMbio strains and subsequently transferred to GFP tagged *Pseudomonas aeruginosa* (PA14-GFP) for 40 hours. **B.** Quantification of GFP fluorescence of worms in A. Note significant protection (*, p<0.05, t-test) offered by *Sphingobacterium* and *Chryseobacterium*. Shown are median and interquartile of N=8-12/group. A similar analysis in worms exposed to PA14-GFP for 36 hours identified also *Pantoea* as significantly protective (not shown).

## References

1. Sommer, F. & Bäckhed, F. The gut microbiota-masters of host development and physiology. Nat. Rev. Microbiol. 11, 227–238 (2013).

2. Morais, L. H., Schreiber, H. L. & Mazmanian, S. K. The gut microbiota–brain axis in behaviour and brain disorders. Nat. Rev. Microbiol. 19, 241–255 (2021).

3. Fan, Y. & Pedersen, O. Gut microbiota in human metabolic health and disease. Nat. Rev. Microbiol. 19, 55–71 (2021).

4. Belkaid, Y. & Harrison, O. J. Homeostatic Immunity and the Microbiota. Immunity 46, 562–576 (2017).

5. Kruger Ben Shabat, S., et al. Specific microbiome-dependent mechanisms underlie the energy harvest efficiency of ruminants. ISME J. 10, 2958–2972 (2016).

6. Rosengaus, R. B., Zecher, C. N., Schultheis, K. F., Brucker, R. M. & Bordenstein, S. R. Disruption of the termite gut microbiota and its prolonged consequences for fitness. Appl. Environ. Microbiol. 77, 4303–4312 (2011).

7. Shukla, S. P., Sanders, J. G., Byrne, M. J. & Pierce, N. E. Gut microbiota of dung beetles correspond to dietary specializations of adults and larvae. Mol. Ecol. 25, 6092–6106 (2016).

8. Gilbert, J. A. et al. Current understanding of the human microbiome. Nat. Med. 24, 392– 400 (2018).

9. Moran, N. A., McCutcheon, J. P. & Nakabachi, A. Genomics and evolution of heritable bacterial symbionts. Annu. Rev. Genet. 42, 165–190 (2008).

10. Feyereisen, R. Arthropod CYPomes illustrate the tempo and mode in P450 evolution. Biochim. Biophys. Acta - Proteins Proteomics 1814, 19–28 (2011).

11. Ceja-Navarro, J. A. et al. Gut microbiota mediate caffeine detoxification in the primary insect pest of coffee. Nat Commun 6, 7618 (2015).

12. Kohl, K. D., Weiss, R. B., Cox, J., Dale, C. & Denise Dearing, M. Gut microbes of mammalian herbivores facilitate intake of plant toxins. Ecol. Lett. 17, 1238–1246 (2014).

13. Shukla, S. P. & Beran, F. Gut microbiota degrades toxic isothiocyanates in a flea beetle pest. Mol. Ecol. 29, 4692–4705 (2020).

14. Alavanja, M. C. R., Hoppin, J. A. & Kamel, F. Health effects of chronic pesticide exposure: Cancer and neurotoxicity. Annu. Rev. Public Health 25, 155–197 (2004).

15. Kikuchi, Y. et al. Symbiont-mediated insecticide resistance. Proc Natl Acad Sci U S A 109, 8618–8622 (2012).

16. Itoh, H., Tago, K., Hayatsu, M. & Kikuchi, Y. Detoxifying symbiosis: Microbe-mediated detoxification of phytotoxins and pesticides in insects. Nat. Prod. Rep. 35, 434–454 (2018).

17. Douglas, A. E. Simple animal models for microbiome research. Nat. Rev. Microbiol. 17, (2019).

18. Shapira, M. Host–microbiota interactions in Caenorhabditis elegans and their significance. Curr. Opin. Microbiol. 38, (2017).

19. Berg, M. et al. Assembly of the Caenorhabditis elegans gut microbiota from diverse soil microbial environments. ISME J. 10, 1998–2009 (2016).

20. Berg, M., Zhou, X. Y. & Shapira, M. Host-Specific Functional Significance of Caenorhabditis Gut Commensals. Front. Microbiol. 7, 1622 (2016).

21. Samuel, B. S., Rowedder, H., Braendle, C., Félix, M.-A. & Ruvkun, G. Caenorhabditis elegans responses to bacteria from its natural habitats. Proc. Natl. Acad. Sci. 113, E3941–E3949 (2016).

22. Berg, M. et al. TGFβ/BMP immune signaling affects abundance and function of C. elegans gut commensals. Nat. Commun. 10, (2019).

23. Ortiz, A., Vega, N. M., Ratzke, C. & Gore, J. Interspecies bacterial competition regulates community assembly in the C. elegans intestine. ISME J. 15, 2131–2145 (2021).

24. Giordano-Santini, R. et al. An antibiotic selection marker for nematode transgenesis. Nat. Methods 7, 721–723 (2010).

25. Ley, R. E. et al. Obesity alters gut microbial ecology. Proc. Natl. Acad. Sci. U. S. A. 102, 11070–11075 (2005).

26. Degruttola, A. K., Low, D., Mizoguchi, A. & Mizoguchi, E. Current understanding of dysbiosis in disease in human and animal models. Inflamm. Bowel Dis. 22, 1137–1150 (2016).

27. Zhang, F. et al. Caenorhabditis elegans as a model for microbiome research. Front. Microbiol. 8, 485 (2017).

28. Dirksen, P. et al. CeMbio - The Caenorhabditis elegans microbiome resource. G3 Genes, Genomes, Genet. 10, 3025–3039 (2020).

29. Wright, G. D. & Thompson, P. R. Aminoglycoside phosphotransferases: proteins, structure, and mechanism. Front. Biosci. 4, D9—21 (1999).

30. Pérez-Carrascal, O. M. et al. Host Preference of Beneficial Commensals in a Microbially-Diverse Environment. Front. Cell. Infect. Microbiol. 12, 1–9 (2022).

31. Melo, J. A. & Ruvkun, G. Inactivation of conserved C. elegans genes engages pathogen- and xenobiotic-associated defenses. Cell 149, 452–466 (2012).

32. Zhang, Z., Liu, L., Twumasi-Boateng, K., Block, D. H. S. & Shapira, M. FOS-1 functions as a transcriptional activator downstream of the C. elegans JNK homolog KGB-1. Cell. Signal. 30, 1–8 (2017).

33. Zhang, Y. et al. Impacts of farmland application of antibiotic-contaminated manures on the occurrence of antibiotic residues and antibiotic resistance genes in soil: A meta-analysis study. Chemosphere 300, 134529 (2022).

34. Cai, M. et al. Occurrence and temporal variation of antibiotics and antibiotic resistance genes in hospital inpatient department wastewater: Impacts of daily schedule of inpatients and wastewater treatment process. Chemosphere 292, 133405 (2022).

35. Stockwell, V. O. & Duffy, B. Use of antibiotics in plant agriculture Fire blight : the primary use of antibiotics on plants activity and mechanisms of resistance in Erwinia amylovora. Rev. sci. tech. Off. int. Epiz. 31, 199–210 (2012).

36. Farkas, A., et al. Molecular Typing Reveals Environmental Dispersion of Antibiotic-Resistant Enterococci under Anthropogenic Pressure. Antibiotics 11, (2022).

37. Ullmann, I. F. et al. Detection of aminoglycoside resistant bacteria in sludge samples from Norwegian drinking water treatment plants. Front. Microbiol. 10, 1–12 (2019).

38. Lester, C. H., Frimodt-Moller, N. & Hammerum, A. M. Conjugal transfer of aminoglycoside and macrolide resistance between Enterococcus faecium isolates in the intestine of streptomycin-treated mice. FEMS Microbiol. Lett. 235, 385–391 (2004).

39. Vitali, L. A., Petrelli, D., Lamikanra, A., Prenna, M. & Akinkunmi, E. O. Diversity of antibiotic resistance genes and staphylococcal cassette chromosome mec elements in faecal isolates of coagulase-negative staphylococci from Nigeria. BMC Microbiol. 14, (2014).

40. Twumasi-Boateng, K. et al. An age-dependent reversal in the protective capacities of JNK signaling shortens Caenorhabditis elegans lifespan. Aging Cell 11, 659–667 (2012).

41. Vind, A. C., Genzor, A. V. & Bekker-Jensen, S. Ribosomal stress-surveillance: Three pathways is a magic number. Nucleic Acids Res. 48, 10648–10661 (2020).

42. Hollos, P., Marchisella, F. & Coffey, E. T. JNK Regulation of Depression and Anxiety. Brain Plast. 3, 145–155 (2018).

43. Lee, J. Y., Tsolis, R. M. & Bäumler, A. J. The microbiome and gut homeostasis. Science (80-.). 377, (2022).

44. Vijay-kumar, M. et al. Metabolic syndrome and altered gut microbiota in mice lacking Toll-like receptor 5. Science (80-.). 344, 228–232 (2010).

45. Ihara, S. et al. TGF-β Signaling in Dendritic Cells Governs Colonic Homeostasis by Controlling Epithelial Differentiation and the Luminal Microbiota. J. Immunol. 196, 4603– 4613 (2016).

46. Xie, K. et al. Dietary S. maltophilia induces supersized lipid droplets by enhancing lipogenesis and ER-LD contacts in C. elegans. Gut Microbes 14, (2022).

47. Dabke, K., Hendrick, G. & Devkota, S. The gut microbiome and metabolic syndrome. J. Clin. Invest. 129, 4050–4057 (2019).

48. Turnbaugh, P. J. et al. An obesity-associated gut microbiome with increased capacity for energy harvest. Nature 444, 1027–1031 (2006).

49. Soen, Y., Knafo, M. & Elgart, M. A principle of organization which facilitates broad Lamarckian-like adaptations by improvisation. Biol. Direct 10, 68 (2015).

50. Bolyen, E. et al. Reproducible, interactive, scalable and extensible microbiome data science using QIIME 2. Nat. Biotechnol. 37, 852–857 (2019).

51. Edgar, R. C. Search and clustering orders of magnitude faster than BLAST. Bioinformatics 26, 2460–2461 (2010).

52. Shapira, M. & Tan, M.-W. Genetic analysis of Caenorhabditis elegans innate immunity. Methods Mol. Biol. 415, 429–442 (2008).

53. Escorcia, W., Ruter, D. L., Nhan, J. & Curran, S. P. Quantification of lipid abundance and evaluation of lipid distribution in Caenorhabditis elegans by nile red and oil red o staining. J. Vis. Exp. 2018, 1–6 (2018).

